# A fully sequenced collection of homozygous EMS mutants for forward and reverse genetic screens in *Arabidopsis thaliana*

**DOI:** 10.1101/2023.10.26.564234

**Authors:** Sébastien Carrère, Jean-Marc Routaboul, Pauline Savourat, Caroline Bellenot, Hernán López, Thomas Quiroz Monnens, Anthony Ricou, Christine Camilleri, Patrick Laufs, Raphael Mercier, Laurent D. Noël

**Affiliations:** LIPME, Université de Toulouse, INRAE/CNRS UMR 0441/2598, Castanet-Tolosan, France; Université Paris-Saclay, INRAE, AgroParisTech, Institut Jean-Pierre Bourgin (IJPB), 78000, Versailles, France; Department of Chromosome Biology, Max Planck Institute for plant breeding research, Carl-von-Linné-Weg 10, Cologne, Germany

## Abstract

Genetic screens are powerful tools for biological research and are one of the reasons for the success of the thale cress *Arabidopsis thaliana* as a model species. Here, we describe the whole-genome sequencing of 871 Arabidopsis lines from the Homozygous EMS Mutant (HEM) collection as a novel resource for forward and reverse genetics. With an average 576 high-confidence mutations per HEM line, over three independent mutations altering protein sequence are found on average per gene in the collection. Pilot reverse genetics experiments on reproductive, developmental and physiological traits confirmed the efficacy of the tool for identifying both null and knockdown alleles. The possibility of conducting subtle repeated phenotyping of HEM lines and the immediate availability of the mutations will empower forward genetic approaches. The sequence resource is searchable with the ATHEM web interface (https://lipm-browsers.toulouse.inra.fr/pub/ATHEM/), and the biological material is distributed by the Versailles Arabidopsis Stock Center.

## Introduction

Model organisms are typically characterised by relative genetic simplicity (gene number, genome size), short reproductive cycle, abundant offspring, amenability for genetic manipulation, small size and straightforward growth conditions. *Arabidopsis thaliana* is a wild Brassicaceae that meets these criteria and has been the main plant model for basic research programs since the 90s^1^. Thousands of forward genetic screens have been conducted worldwide using this plant, Columbia-0 (Col-0) being the most commonly used accession. Such screens commonly require the phenotyping of tens of thousands of individuals, thus limiting our capacity to conduct labour-intensive or expensive phenotypic characterizations. With more than 32,000 genes annotated in the Col-0 reference genome, using homozygous T-DNA insertion mutants also represents an important workload. Besides, such mutants are mostly null loss-of-function mutants, which limits the range of induced genetic variations and hinders the isolation of mutants in essential genes whose analysis would benefit from hypomorphic or conditional alleles.

In order to circumvent some of these limitations, a set of 897 homozygous EMS mutant (HEM) lines were produced by single seed descent or haploid doubling in the Col-0 accession^2^. The whole-genome sequence analysis of 47 HEM lines has previously shown that each line carried ca. 700 homozygous mutations, 28% of which affect a protein sequence. By construction, most of the mutations in the HEM lines are fixed, which means that phenotypes can be replicated and multiple traits can be explored in these immortalised lines. A previous forward genetic screen looking for meiotic defects identified 43 lines, 21 of which carried mutations in genes previously identified to have a key role in meiosis. Several allelic series were also found, suggesting that sequencing the whole HEM library could facilitate the identification of the causal genes.

In this study, through whole-genome sequencing of 871 HEM lines, we describe the complete set of mutations present in the collection. We experimentally validated a selected subset of detected mutations, at both the molecular and phenotypic levels. To help the community exploit this new resource, we have constructed a web interface to facilitate forward and reverse genetic approaches with the HEM collection.

## Results

### Sequencing of HEM lines and detection of EMS-induced mutations

DNA was extracted from pooled leaf samples originating from five seedlings of each of the 897 HEM lines and 71 wild-type Col-0 controls. Genomic DNA was sequenced on three NovaSeq6000 lanes (Illumina) yielding 2.43 Tb of sequences (8.1 billions paired-end reads of 2 x 150bp). Quality-filtered reads (e.g., adapter trimming, removal of duplicates) from each HEM line were mapped to the reference genome using relaxed criteria in order to minimise the chance of missing EMS-induced polymorphisms (see Material and Methods for detailed parameters). Considering a genome size of 119 Mb, the effective median coverage is 10.06 per HEM line (Fig. 1a). This analysis identified 433,918 putative polymorphic sites in the 871 HEM lines. Twenty-six HEM lines were not retained for further analyses due to poor sequence coverage or aberrant patterns of polymorphisms. As expected for an EMS mutagenesis, 96% of the polymorphisms corresponded to SNPs, 97% of which were transitions (G->A or C->T). The other observed polymorphisms resulted from small InDels (1% insertions, 3% deletions). Considering only transition mutations falling into genes (including UTRs and introns), 85% (251,285) were predicted to be homozygous (refer to Materials and Methods).

**Fig. 1:**
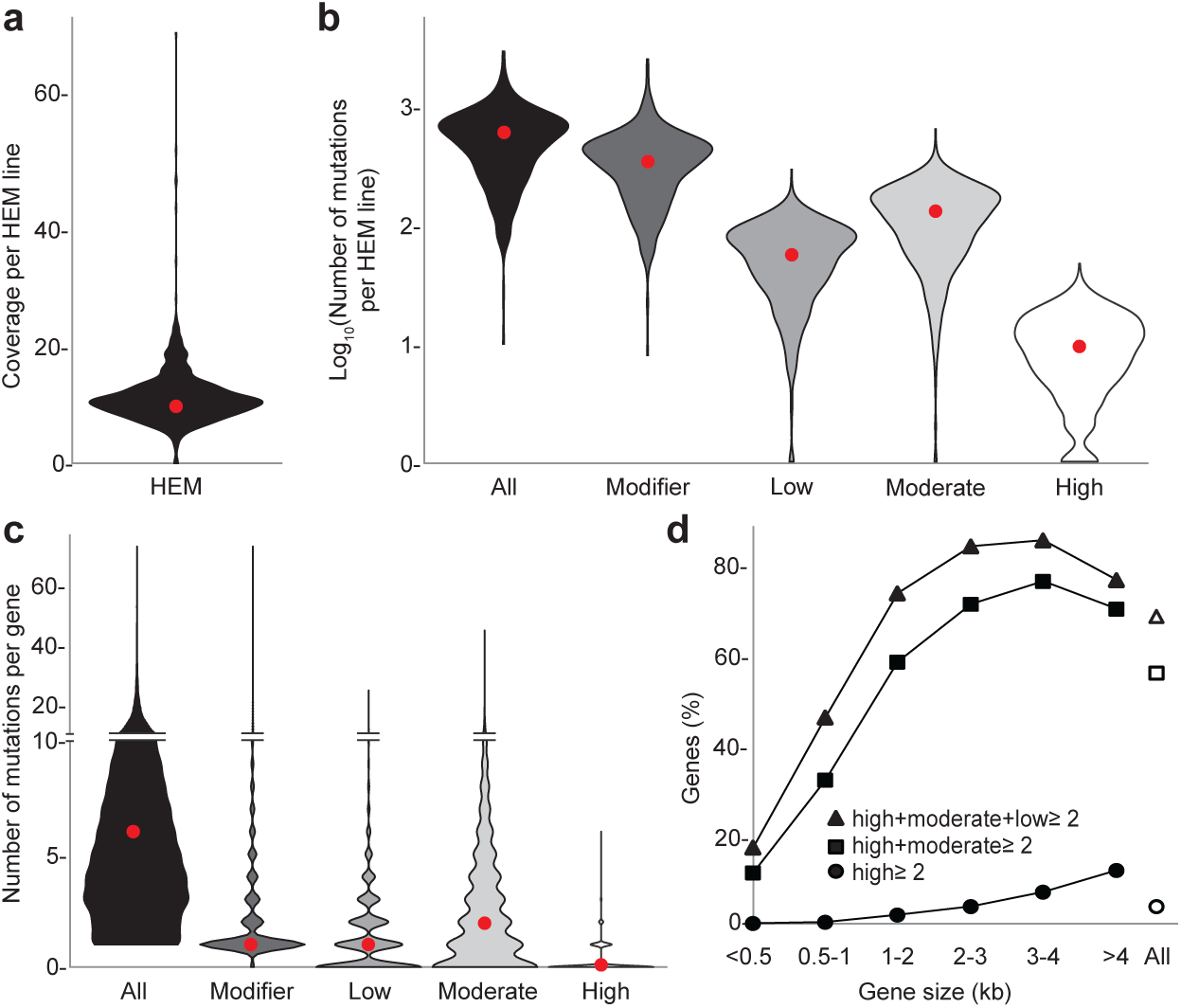
Features of the genome sequences of 871 HEM lines. **a**, violin plot representation of the distribution of mean coverage of sequencing per HEM line. **b**, violin plot representation of the distribution of the number of mutations per line and per predicted impact of mutation. **c**, violin plot representation of the distribution of the number of mutations per gene and per predicted impact of mutation. The maximum width of each violin was made equal to increase the readability, so that the surface area of each violin is not proportional to their sample size. **d**, Proportion of genes of given sizes for which at least two mutations of the impact categories high (dot), high+moderate (square) or high+moderate+low (triangle) can be found in the HEM collection. Open symbols correspond to all genes independent of their size. Red dots represent medians. Impact of mutations: high, premature stop codon, splicing alteration or frameshift mutation; moderate, nonsynonymous substitution; low, synonymous substitution; modifier, unpredictable effect of mutations in non-coding regions of a gene or in non-coding genes.

### Prediction of the impact of HEM polymorphisms on gene function

On average, 576 mutations were detected per HEM line, 135 (23%) of which are predicted to have a high or moderate impact on gene function (Table 1, Fig. 1b) using the SnpEff software (https://pcingola.github.io/SnpEff/se_inputoutput/#effect-prediction-details). High-impact mutations include premature stop codons, frameshift mutations or splicing alterations that likely result in protein truncation. Moderate-impact mutations correspond to missense mutations or inframe deletions that might affect protein function. Low or modifier mutations correspond to synonymous mutations or mutations in non-coding regions, respectively, and together represent 77% of the polymorphisms. All of the 32,723 genes of Arabidopsis accession Col-0 harbour one or more detected mutations, with an average of 3.25 polymorphisms per kilobase. Importantly, 20% and 73% of the genes harbour at least one high-impact or high/moderate-impact mutation, respectively (Fig. 1c). The 8,700 (27%) genes without high– and moderate-impact mutations might be small and/or essential for viability or reproduction. Indeed, we observed a positive correlation between gene size and mutation frequency (Fig. 1d). In conclusion, it is expected that at least two high– or moderate-impact mutations can be identified for 57% of Arabidopsis genes (Fig. 1d), thus contributing to a high likelihood of identifying allelic series in the HEM resource. This collection is thus suitable for use in both forward and reverse genetic screens.

**Table 1:**
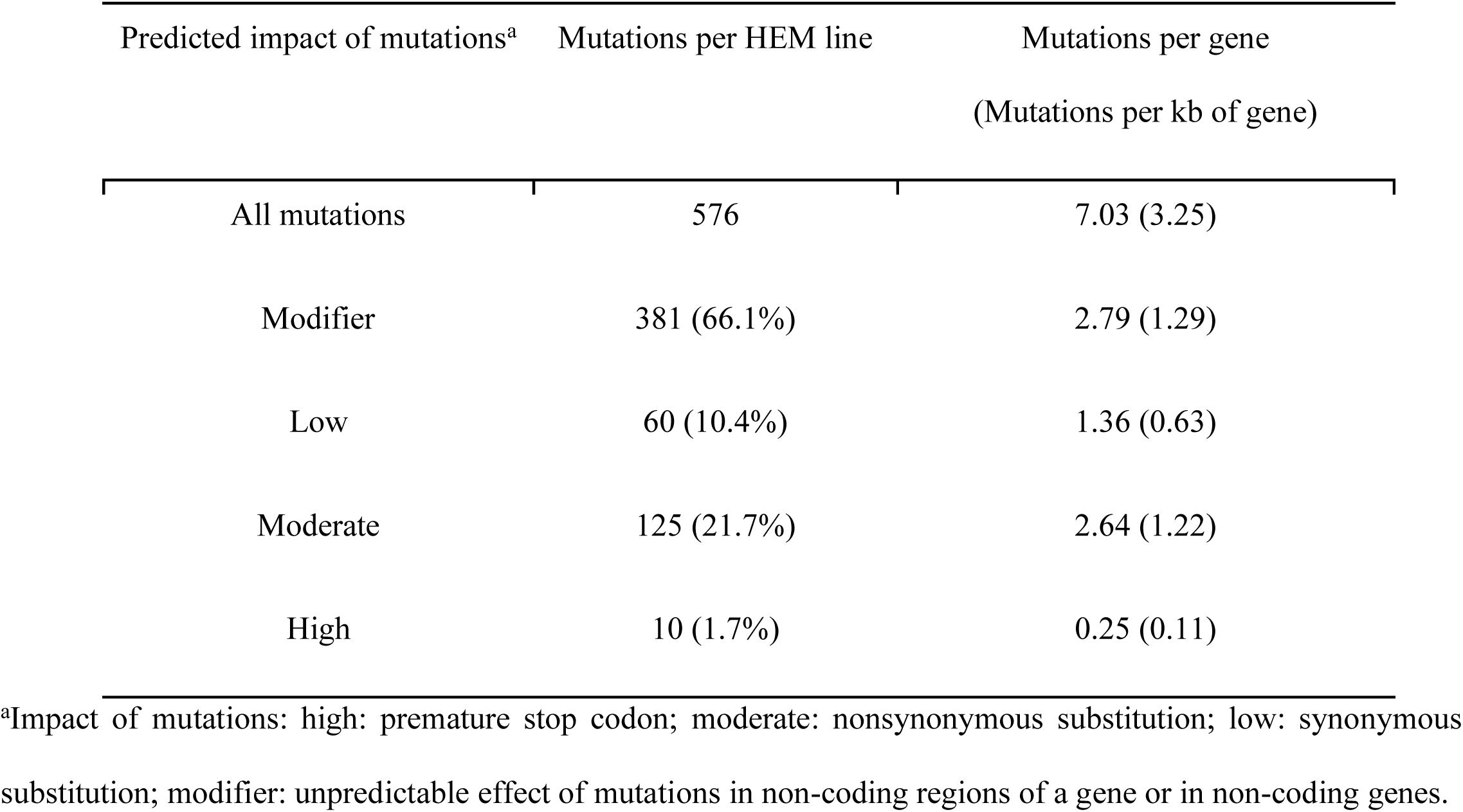
Nature and mean number of mutations detected in 871 HEM lines and 32,723 genes.

| Predicted impact of mutations<sup>a</sup> | Mutations per HEM line | Mutations per gene<br>(Mutations per kb of gene) |
| --- | --- | --- |
| All mutations | 576 | 7.03 (3.25) |
| Modifier | 381 (66.1%) | 2.79 (1.29) |
| Low | 60 (10.4%) | 1.36 (0.63) |
| Moderate | 125 (21.7%) | 2.64 (1.22) |
| High | 10 (1.7%) | 0.25 (0.11) |

<sup>a</sup>Impact of mutations: high: premature stop codon; moderate: nonsynonymous substitution; low: synonymous substitution; modifier: unpredictable effect of mutations in non-coding regions of a gene or in non-coding genes.

### Web-based search interface to mine HEM SNP repertoire

A web-based interface named ATHEM was created to visualise sequence alignment results per HEM line, per gene, per genomic region and per impact of mutation (https://lipm-browsers.toulouse.inra.fr/pub/ATHEM/). This site also offers a user-friendly searchable tool, including a genome browser, to allow mining of the resource for forward and reverse genetics applications. The user can evaluate the quality of the predicted polymorphisms and the homozygosity state based on sequence alignments using the genome browser.

### Mutations inferred from the HEM database sequences reliably identify homozygous mutants for reverse genetic approaches

We conducted a number of quality controls which first included the comparison of the polymorphisms detected here in 47 sequenced lines with those previously identified in their progenitors^2^. The analysis showed that the closest hit for each progenitor was indeed its expected HEM descendant. In a previous forward genetic screen for meiotic defects conducted on the HEM collection^2^, causal mutations were identified in 18 lines re-sequenced here. Among these 18 mutations, 11 were found in the HEM database in the expected lines. The seven remaining mutations are not found in the HEM database. This could be explained by genetic segregation as the initial forward screen ^2^ was performed in an earlier generation than the mutant plants sequenced here.

Next, to further investigate the quality and potential for functional analysis of this population, we conducted reverse genetic screens with five functionally well-characterised genes (Table 2). These included genes involved in flavonoid production in seeds, meiosis or shoot and flower development. A first reverse genetic screen was undertaken for the *CHALCONE SYNTHASE (TT4)* gene and the *TRANSPARENT TESTA 2 (TT2)* gene encoding a R2R3 MYB domain transcription factor. These genes are both key determinants in the accumulation of flavonoids, including proanthocyanidins, that are responsible for the brown colour of mature seeds. Four and three HEM lines were predicted to have high– or moderate-impact mutations in *TT4* and *TT2*, respectively. Sanger sequencing of five individual plants per line showed that all seven HEM lines carried their expected mutation in *TT2* or *TT4*. Six mutations were homozygous and one was segregating (ES1M5S03056) (Table 2), matching the Illumina sequencing. The three lines predicted to have a high impact (ES1M5S02007 and EH1S1B627 for *TT2*; ES1M5S10055 for *TT4*) and a single one predicted to have moderate impact on *TT4* (EH1S1B670) displayed a yellow colour indicative of a lack of proanthocyanidin accumulation in seeds (Fig. 2a), as expected for a null mutation in this metabolic pathway.

**Fig. 2:**
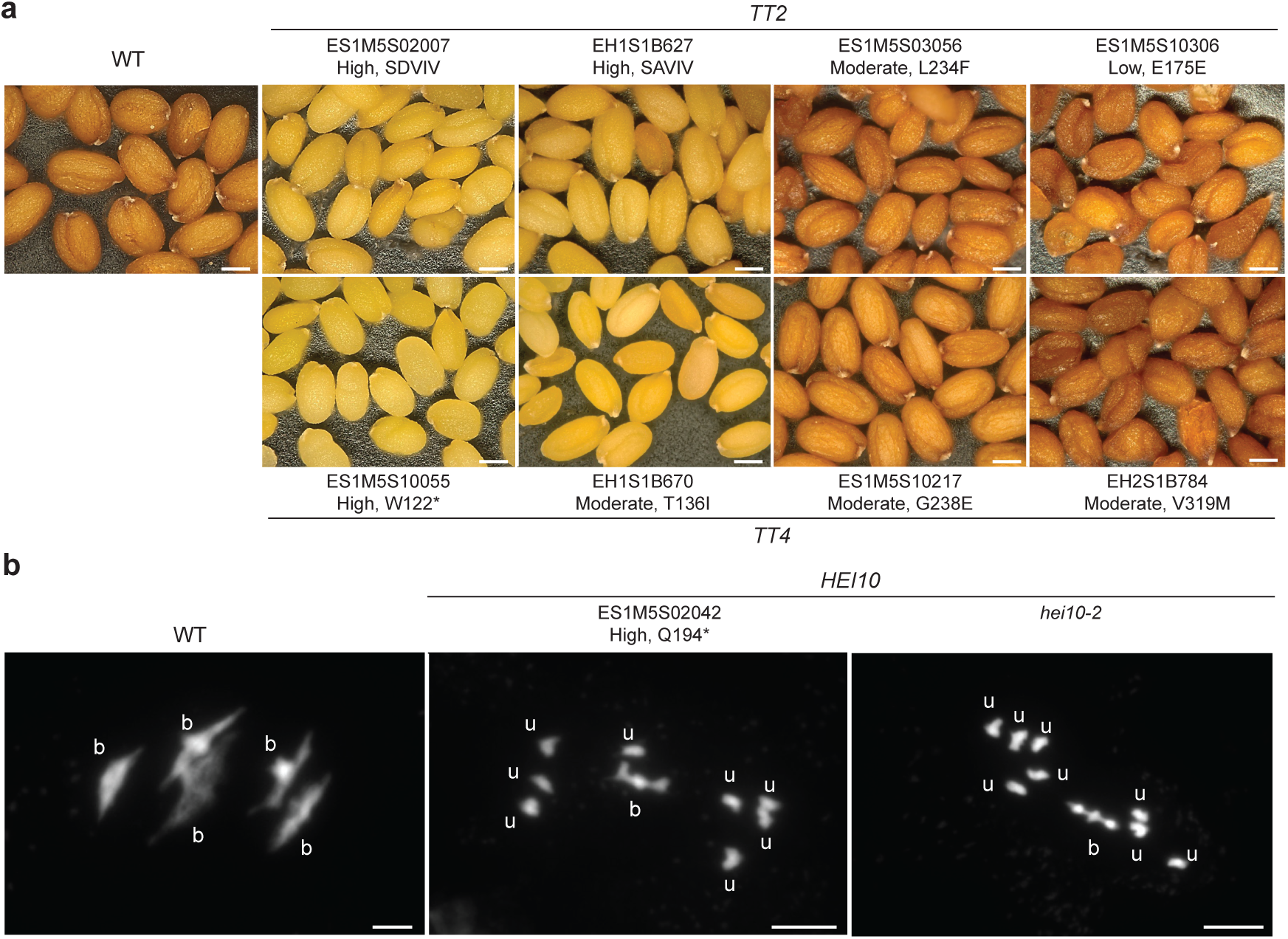
Selected mutant phenotypes identified by reverse genetics in HEM lines. **a**, Seeds of HEM lines mutant for *TT2* and *TT4* genes controlling flavonoid accumulation. HEM line and predicted functional impact of the mutation on TT gene function: High, moderate, SDVIV (Splice donor variation and intron variant); SAVIV (Splice acceptor variation and intron variant). WT, wild-type accession Col-0. Impact of the mutation on protein sequence is indicated. Scale: 0.2 mm. **b**, Chromosome spreads of wild-type (WT), *hei10^Q^*^194^*\** (ES1M5S02042) and *hei10-2* male meiocytes at metaphase I. Wild type shows five bivalents (b) whereas in the representative spreads of *hei10^Q^*^194^*\** and *hei10-2*, the meiocytes showed one bivalent and four pairs of univalent (u) chromosomes. Scale: 5 µm.

**Table 2:** Phenotypic characterization of HEM mutant phenotypes by reverse genetics.

| Gene | Null mutant phenotype | Gene size (kb) | Predicted lines | Observed | Homozygous | Mutant phenotype | Reference |
| --- | --- | --- | --- | --- | --- | --- | --- |
|  |  | 5’->3’ UTR | with H/Mr/L/Mf | H/Mr/L/Mf | H/Mr/L/Mf SNPs<sup>a,b,c</sup> | observed for |  |
|  |  |  | SNPs<sup>a,b</sup> | SNPs<sup>a,b</sup> |  | H/Mr/L/Mf mutants<sup>a,b</sup> |  |
| *TT2* (At5g35550) | Transparent testa seeds | 1.1 | 2/1/3/6 | 2/1/NT/NT | 2/0/NT/NT | 2/-/-/- | <sup>25</sup> |
| *TT4* (At5g13930) | Transparent testa seeds | 1.6 | 1/3/3/3 | 1/3/NT/NT | 1/3/NT/NT | 1/1/-/- | <sup>26</sup> |
| *CUC1* (At3g15170) | Cup-shaped cotyledon | 1.6 | -/7/-/3 | -/7/-/NT | -/7/-/NT | -/7/-/NT | <sup>27</sup> |
| *CUC2* (At5g53950) | Cup-shaped cotyledon | 1.9 | -/5/5/4 | -/5/1<sup>d</sup>/NT | -/5/1<sup>d</sup>/NT | -/4/1<sup>d</sup>/NT | <sup>4</sup> |
| *HEI10* (At1G53490) | Meiotic Crossover<br>formation | 3.6 | 1/1/3/14 | 1/1<sup>e</sup>/NT/NT | 0/0/-/- | 1/NT/-/- | <sup>3</sup> |

<sup>a</sup> Predicted impact of mutations: high (H), moderate (Mr), low (L) and modifier (Mf).
<sup>b</sup> NT: not tested
<sup>c</sup> At least one homozygous mutant plant identified in five plants genotyped
<sup>d</sup> A single low-impact mutant (ES1M5S03057) was analysed for *CUC2*.
<sup>e</sup> A single moderate-impact mutant (ES1M5S10109) was analysed for *HEI10*.

HEI10 is an evolutionarily conserved RING finger-containing protein involved in the formation of meiotic crossovers^3^. Meiotic crossovers shuffle genetic information and create physical links between homologous chromosomes – chiasmata – which are essential for balanced chromosome segregation. In the absence of a functional HEI10, crossover (CO) formation is strongly reduced, unconnected chromosomes (univalents) segregate erratically at meiosis I, and fertility is impaired. We identified two mutations in the HEM lines that modify the *HEI10* coding region (Table 2). One mutation with predicted high impact in the ES1M5S02042 line (C580T) introduces a premature stop at codon 194 (Q194*) and is predicted to result in the production of a HEI10 protein truncated at its C-terminal unstructured region, based on AlphaFold modelling. The second mutation (C800T) present in the ES1M5S10109 line is predicted to have a moderate impact (P267L). We confirmed that both mutations were present and segregated in the ES1M5S02042 and ES1M5S10109 lines, corroborating the whole-genome sequencing data (Table 2 and Supplementary Table S1). Plants homozygous for the *hei10*^Q194*^ mutation showed strongly reduced fertility, as assessed by visual examination of fruit length. Meiotic chromosome spreads revealed the presence of univalents, phenocopying the previously described *hei10-2* mutant (Fig. 2b). This shows that the *hei10*^Q194*^ mutation disrupts *HEI10* function and indicates that the C-terminal unstructured region of HEI10 is important for its function in CO formation.

We finally searched for mutants in the *CUP-SHAPED COTYLEDON1* (*CUC1*) or *CUC2* genes which are required for boundary domain specification in the aerial organs. Mutants in these genes show multiple phenotypes including fusion between organs such as cotyledons and sepals and reduced leaf serration for *cuc2*^4,5^. Seven and five HEM lines were predicted to have high– or moderate-impact mutations in *CUC1* and *CUC2*, respectively. Sequencing of *CUC1* or *CUC2* in eight individual plants per HEM line showed that most were homozygous mutants as inferred from genome sequences. The only exceptions were the ES1M5S03067 and ES1M5S11077 lines in which, among the eight plants investigated, a single wild-type plant for the *CUC1* and *CUC2* genes was identified, respectively. In the absence of heterozygous plants in the same HEM lines, we hypothesise that those plants wild-type for *CUC1* or *CUC2* may result from occasional seed contamination that occurred during seed collection or after sowing. Mutations affected both conserved and nonconserved amino acid residues of CUC proteins (Supplementary Fig. S1). All seven *cuc1* and four out of five *cuc2* candidate moderate-impact mutants showed fused cotyledons and/or sepals (Table 2, Supplementary Table S1, Fig. 3a,b). The *cuc2*^P59L^ *and cuc2*^G196D^ mutants also showed reduced leaf serration (Fig. 3c). These phenotypes were indicative of impaired CUC1 and CUC2 function. To further test if the phenotypes were due to mutations in *CUC1* or *CUC2* genes, we performed allelism tests. F1 plants of a cross between either one of the two of the strongest *cuc1* HEM mutants based on the sepal fusion phenotype (*cuc1*^E75K^ or *cuc1*^G120R^) with the strong mutants *cuc1-13* showed fused sepals. Similarly, F1 plants resulting from the cross of either *cuc2*^P59L^ or *cuc2*^G196D^ with the strong mutant *cuc2-1* showed fused sepals and reduced leaf serration (Fig. 3c). Together, this confirmed that the phenotypes observed in the HEM lines were due to mutation of *CUC1* or *CUC2*. Double *cuc1 cuc2* mutants show strong cotyledon fusion defects and form cup-shaped cotyledons^4^. Similar cup-shaped cotyledon seedlings were formed in the F2s between *cuc2-1* and *cuc1*^E75K^ or *cuc1*^G120R^, suggesting that these two *CUC1* alleles severely affected its function (Fig. 3a). In contrast, no cup-shaped cotyledon phenotype was observed in the F2 population between *cuc1-13* and the two novel *cuc2*^P59L^ and *cuc2*^G196D^ alleles, suggesting that these are hypomorphic alleles, consistent with the limited suppression of their leaf serration compared to the smooth leaves of the strong *cuc2-1* allele (Fig. 3c)^6^. Altogether, these observations indicated that both strong and hypomorphic alleles of *CUC1* and *CUC2* could be identified in the HEM collection.

**Fig. 3:**
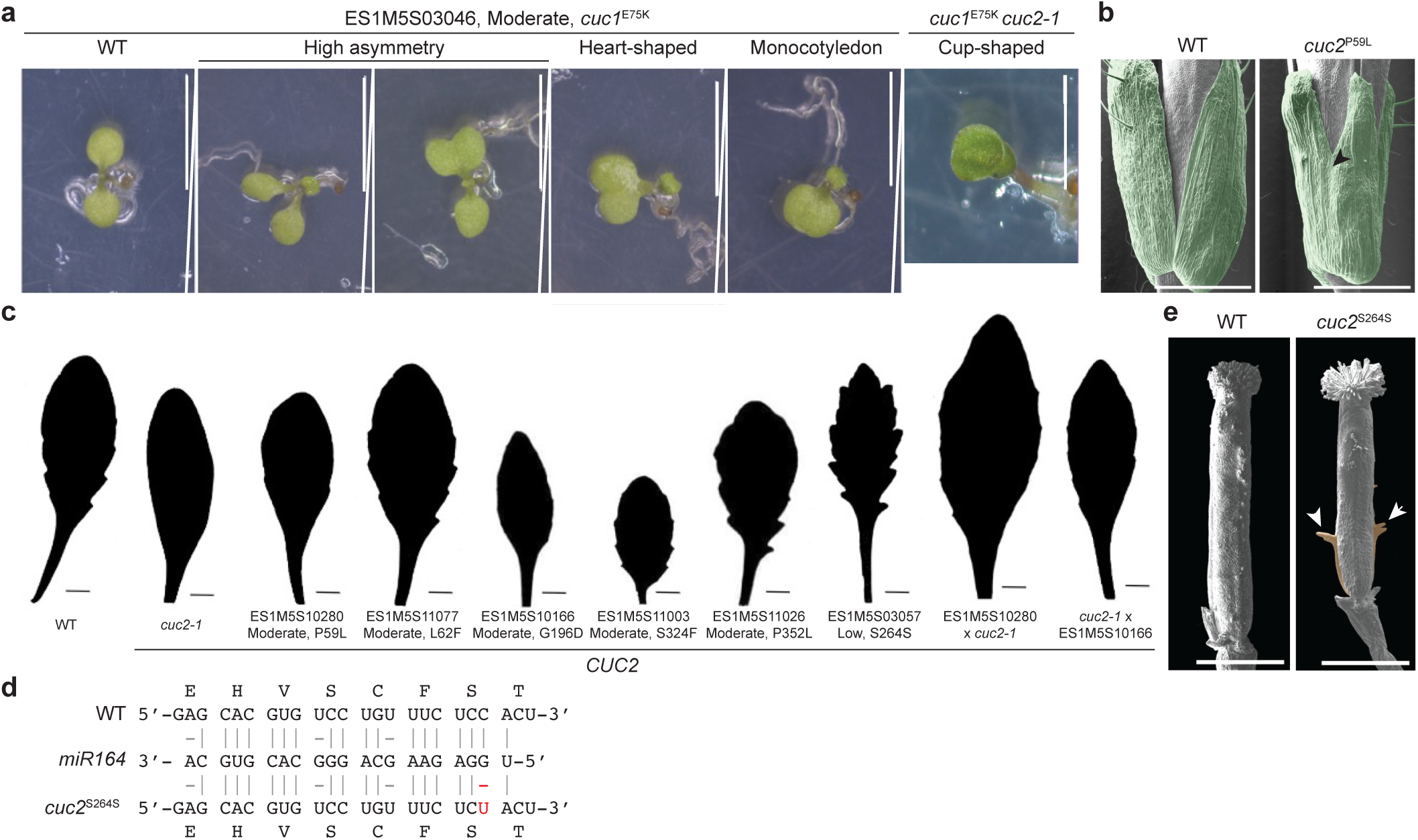
Phenotypic description of *cuc1* and *cuc2* mutants identified in HEM lines. **a**, Cotyledon phenotypes ranging from no fusion (WT, wild-type Col-0) to monocotyledon or cup-shaped cotyledon observed in the ES1M5S03046 HEM line carrying the *cuc1*^E75K^ mutation and the *cuc1*^E75K^*cuc2-1* double mutant. Scale: 4 mm. **b**, Sepal fusion (Arrohead) observed by scanning electron microscopy in the ES1M5S10280 HEM line carrying the *cuc2*^P59L^ mutation compared to wild-type Col-0 plants. False colours were used to visualise sepals (green). Scale: 1 mm. **c**, Representative shape of the 8^th^ leaf of the rosette of HEM lines carrying mutations in *CUC2* and their F1 progenies of a cross with the strong *cuc2-1* mutant. Scale: 10 mm. **d**, Alignment of RNA sequences for wild-type *CUC2*, *miR164* and *CUC2* allele from HEM line ES1M5S03057 (*cuc2*^S264S^). Polymorphic base is highlighted in red. **e**, Outgrowths (orange) observed on the pistil of HEM line ES1M5S03057 (*cuc2*^S264S^) compared to the wild type are indicated by arrowheads. Scale bar is 1mm.

Our results showed that the mutations inferred from the HEM database were reliably confirmed, that the corresponding homozygous mutants could be easily retrieved in the HEM collection and that both strong and hypomorphic mutants for those genes of interest could be identified.

### A unique cuc2 mutation in the miR164 binding sequence important for the repression of CUC2 expression

While identifying mutations that affect protein sequence may be the primary use of the HEM collection, other mutations classified as Low or Modifier may nevertheless be useful and yield informative mutant phenotypes. For instance, we identified the ES1M5S03057 HEM line as containing a synonymous point mutation in *CUC2* (S264S) inside the known binding site for *miR164* microRNA, a negative regulator of *CUC1* and *CUC2* expression (Fig 3d)^6,7^. Interestingly, the *cuc2*^S264S^ line showed increased leaf serration (leaf dissection index was 1.30 on the 6^th^ rosette leaf compared to 1.23 for wild type, n>14), and ectopic structures developed along the pistil (Fig. 3e). These phenotypes are very reminiscent of the *cuc2*-1D mutant, which was demonstrated to disrupt the *miR164* binding site resulting in increased *CUC2* expression^8^. This suggested that the regulation of the endogenous *CUC2* gene by miR164 was compromised in *cuc2*^S264S^. This example illustrates the usefulness of a fully sequenced collection of SNP mutants for the identification of rare alleles affecting regulation of gene expression.

## Discussion

### Using the HEM resource to go from genes to phenotypes, and *vice versa*

Here, we fully sequenced a collection of more than 800 Arabidopsis mutants, thus generating a new genetic resource for forward and reverse genetics that we are making available for the community. This resource is complementary to the numerous insertion mutant collections available for Arabidopsis and parallels those developed for other species such as wheat^9^.

To demonstrate the usefulness of this resource, we first confirmed the reliability of our SNP analysis by Sanger-sequencing of 23 gene fragments from 23 distinct HEM lines. Second, we identified novel alleles in well-characterised reproductive, developmental or physiological processes, including loss-of-function, hypomorphic and gain-of-functions alleles resulting in gene expression misregulation that could not have been found in insertional mutant collections. We have developed a user-friendly web interface that allows for the efficient mining of the resource. With over 500 mutations per line, characterization of given mutants identified in reverse genetic approaches will still require backcrossing and/or complementation. Identification of allelic series within the HEM collection is possible and may be used to rapidly identify candidate causal mutations linked with phenotypic defects. Because the saturation of the mutagenesis still remains limited, one may still proceed to a mapping-by-sequencing strategy for mutants of particular significance without allelic series.

### The HEM resource in the landscape of Arabidopsis mutant collections

Numerous genetic tools are available for conducting reverse genetic screens in Arabidopsis. These include large transposon or T-DNA mutant collections, some of which are homozygous (https://arabidopsis.info/BrowsePage). This allowed the assembly of homozygous mutant collections for specific gene families such as root-expressed LRR-RLK (69 genes)^10^ or disease resistance genes (ARTIC collection, 171 genes)^11^. A number of large TILLING (Targeting Induced Local Lesions IN Genomes) resources produced by EMS mutagenesis in accessions Col-*erecta*, C24 and Landsberg *erecta* were also generated and were used successfully^12–14^, though the trend would be to prefer CRISPR-generated mutants for targeted reverse genetic approaches. TILLING by sequencing relies on high-throughput amplicon sequencing on DNA pools to identify desired mutants in collections^15^. Yet, whole genome sequencing of such TILLING resources is still difficult to consider due to the constant segregation of EMS-induced mutations and the population sizes which usually exceed tens of thousands of individuals. Furthermore, the heterozygous nature of such TILLING populations would render some forward genetic screens particularly tedious. With the high frequency of homozygous mutations, the HEM collection is therefore particularly valuable for reverse genetic approaches. Beyond null alleles that can attribute a function to a gene, specific alleles can provide important insights into the mechanistics or the regulation of important developmental or physiological processes, as exemplified here. The HEM collection thus provides a fully sequenced collection of homozygous mutants whose population size is suitable with both reverse and forward genetic applications.

### The HEM core mutant collection

The “immortal” nature of the HEM collection, a property resulting from the homozygosity of the mutants, will allow the community to conduct large-scale phenotyping on a small core collection of sharable stable mutants. This will allow scientists to conduct screens based on omics methodologies (transcriptomics, metabolomics, epigenetics, proteomics) as well as labour-intensive forward screens (molecular, microscopic, biochemical) that would otherwise not be conducted on larger collections. Furthermore, because the screens will be conducted on the same material, they have the potential to reveal correlations between different traits quantified in independent screens. HEM lines should also identify genes other than those revealed by genome-wide association studies in natural accessions, which may not be polymorphic due to strong natural selection. It will thus be exciting to test in the near future whether such approaches will unveil unexpected correlations between phenotypes and biological processes that would not have been connected otherwise. Last but not least, the EMS-generated HEM mutants are considered as non-GMO organisms which can thus be freely used for *in natura* screens.

## Online Methods

### Plant material

The 897 homozygous EMS mutant (HEM) lines (Col-0 accession) were obtained by either single seed descent or haploid doubling^2^ and are available from the Versailles Arabidopsis stock center (https://publiclines.versailles.inrae.fr/catalogue/hem). Plants were grown in a greenhouse or in a growth chamber on *Jiffy*-7® peat pellets (http://www.jiffypot.com) under short-day conditions (8h light, 100-120µE).

### Sequencing of genomic DNA from the HEM line

Single 3-mm leaf discs were sampled from five seedlings per HEM line, pooled and subjected to DNA extraction and library preparations by the Max Planck Genome Center, as described^16^. Seventy-one wild-type Col-0 plants were included in the sequencing protocol as internal references. Sequencing libraries were multiplexed in three pools for sequencing. Paired-end sequencing (2×150bp) was conducted on NovaSeq6000 (Illumina) in three pools yielding 2.43 Tb (8.1 x 10^9^ paired reads). Libraries with initially low sequencing output were resequenced on Nextseq2000.

### Sequence analyses and SNP identification

Raw sequence reads were processed using a new Nextflow pipeline nf-mutdetect2. This pipeline first trimmed raw reads based on quality scores using trimmomatic software^17^ (parameters: LEADING:20 TRAILING:20 SLIDINGWINDOW:4:20 MINLEN:50). Trimmed reads were mapped onto the reference genome of *A. thaliana* (TAIR10) with bwa-mem^18^ (default parameters), and alignments were filtered to remove duplicates and keep only paired alignments with the samtools suite^19^ (parameters: samtools view –f 0×02 | samtools fixmate –c –r | samtools rmdup | samtools view –b –q 1 –F 4 –F 256). SNPs and small INDELs were called using samtools mpileup (parameters: samtools mpileup –B ––max-depth 100) and varscan (HEM lines parameters: ––min_coverage=3 –– min_reads2=3 ––avg_qual=15 ––var_freq=0.2 ––var_freq_for_hom=0.8 ––pvalue=0.01; Wild-Type lines parameters: ––min_coverage=5 ––min_reads2=4 ––avg_qual=15 ––var_freq=0.2 –– var_freq_for_hom=0.8 ––pvalue=0.01) tools^20^. Polymorphic sites found in HEM lines and in at least three parental Col-0 lines (out of 71 sequenced individuals) were excluded, because these mutations are likely originating from the parent. We also excluded polymorphisms shared between more than four HEM lines, since mutations induced by EMS are expected to be random and distinct between distinct HEM lines. A final filtering step was applied in order to discard variation with low impact predicted by the SNPeff tool^21^ and the TAIR10 genome annotation.

### Setup of a HEM searchable web tool

All informative intermediate data files such as clean alignments and complete variation matrices are provided on a dedicated web site https://lipm-browsers.toulouse.inra.fr/pub/ATHEM/. This website also provides access to summary tables (list of genes with a mutation in each line and list of lines showing a mutation in each *A. thaliana* gene, all classified by predicted impact), statistical summaries and finally a search engine with direct access to pre-filtered variation sites (chromosomal sites with a maximum coverage of 100x). Users can look for genes/lines or chromosomal regions and select only a minimum impact level.

Polymorphic sites can be displayed on a dedicated genome browser providing both clean alignments and genome annotations.

### Phenotyping of HEM mutants for reproductive, developmental and physiological responses

Putative mutant plants were grown in the greenhouse or growth chambers. Genomic DNA was extracted from five to eight individual plants. The region of the target gene containing the mutation was amplified by PCR using the appropriate primers (Supplementary Table S2), and the PCR product was sequenced. The phenotype was observed on the same set of plants. Pictures for HEM lines mutant for *TT2* and *TT4* genes controlling flavonoid accumulation were taken using a Zeiss Axio Zoom V16 with a Plant Neo Fluar Z 1.0X objective and brightfield reflected light. Meiotic chromosome spreads were performed as described^22^. Leaf morphology was determined using MorphoLeaf software^23^. Scanning electron microscopy of sepals was performed as described^24^.

## Data and Software Availability

The sequence datasets generated and analysed during the current study are available in the NCBI Sequence Read Archive (SRA) repository under accession SRP429727. The source code of the pipeline used to analyse these datasets is available at https://github.com/lipme/nf-mutdetect2. HEM seeds are available from the Versailles Arabidopsis Stock Center (https://publiclines.versailles.inrae.fr/catalogue/hem)

## Supporting information

Supplementary TablesS1 and S2

## Acknowledgements

We wish to thank all SIX team members for their sampling effort (LIPME, Toulouse, France; www.xantho.fr) and in particular Carine Gris for the resequencing of the *TT2 and TT4* genes. We would also like to thank the Max Planck Genome Center for library preparation and sequencing (Cologne, Germany).

## Supplemental Tables

**Table S1: Sequence and phenotypic analyses of allelic series identified in the HEM collection for five genes of interest**

**Table S2: Oligonucleotide sequences used in this study**

## Supplemental Figure

**Fig. S1: Protein sequence alignment of CUC2 and CUC1 orthologues**

