## Supplementary TablesS1 and S2 for "A fully sequenced collection of homozygous EMS mutants for forward and reverse genetic screens in *Arabidopsis thaliana*"

**Table S1: Sequence and phenotypic analyses of allelic series identified in the HEM collection for five genes of interest**

| | | Prediction based on sequencing | | | | Confirmation | | | |
|---|---|---|---|---|---|---|---|---|---|
| Gene | HEM line | Impact on gene | Protein change | WT[a] | Mutant[a] | WT | Heterozygous | Mutant | Phenotype |
| *TT2* (At5g35550) | ES1M5S02007 | High | Splicing | 0 | 8 | 0/5 | 0/5 | 5/5 | Yellow seeds due to absence of brown pigment in seed coat (testa). |
| | EH1S1B627 | High | Splicing | 0 | 6 | 0/5 | 0/5 | 5/5 | Yellow seeds due to absence of brown pigment in seed coat (testa). |
| | ES1M5S03056 | Moderate | L234F | 3 | 7 | 2/5 | 2/5 | 1/5 | - |
| | ES1M5S10306 | Low | E175E | 0 | 11 | ND[b] | ND | ND | - |
| *TT4* (At5g13930) | ES1M5S10055 | High | W122* | 0 | 5 | 0/5 | 0/5 | 5/5 | Yellow seeds due to absence of brown pigment in seed coat (testa). |
| | EH1S1B670 | Moderate | T136I | 0 | 15 | 0/5 | 0/5 | 5/5 | Yellow seeds due to absence of brown pigment in seed coat (testa). |
| | ES1M5S10217 | Moderate | G238E | 0 | 10 | 0/5 | 0/5 | 5/5 | - |
| | EH2S1B784 | Moderate | V319M | 0 | 11 | 0/5 | 0/5 | 5/5 | - |
| *CUC1* (At3g15170) | EH3S1B711 | Moderate | L38F | 0 | 18 | 0/8 | 0/8 | 8/8 | Weak cotyledon fusion (0.11%), weak sepal fusion |
| | ES1M5S03067 | Moderate | G72E | 1 | 17 | 1/8 | 0/8 | 7/8 | Weak cotyledon fusion (0.17%) |
| | ES1M5S03046 | Moderate | E75K | 0 | 22 | 0/8 | 0/8 | 8/8 | Strong cotyledon fusion (7.10%), strong sepal fusion |
| | ES1M5S10276 | Moderate | G120R | 0 | 6 | 0/8 | 0/8 | 8/8 | Weak cotyledon fusion (0.37%), strong sepal fusion |
| | ES1M5S06139 | Moderate | G149D | 0 | 10 | 0/8 | 0/8 | 8/8 | Weak cotyledon fusion (0.14%) |

| | Prediction based on sequencing | | | | | Confirmation | | | |
|---|---|---|---|---|---|---|---|---|---|
| | ES1M5S10182 | Moderate | S177N | 0 | 13 | 0/8 | 0/8 | 8/8 | No cotyledon fusion (0%), weak sepal fusion |
| | ES1M5S01031 | Moderate | P234L | 0 | 6 | 0/8 | 0/8 | 8/8 | No cotyledon fusion (0%), weak sepal fusion |
| *CUC2* (At5g53950) | ES1M5S10280 | Moderate | P59L | 0 | 10 | 0/8 | 0/8 | 8/8 | Strong cotyledon fusion (1.71%), strong sepal fusion, reduced leaf serration |
| | ES1M5S11077 | Moderate | L62F | 1 | 10 | 1/8 | 0/8 | 7/8 | - |
| | ES1M5S10166 | Moderate | G196D | 0 | 8 | 0/8 | 0/8 | 8/8 | No cotyledon fusion (0%), strong sepal fusion, reduced leaf serration |
| | ES1M5S11003 | Moderate | S324F | 0 | 5 | 0/8 | 0/8 | 8/8 | Weak cotyledon fusion (0.11%) |
| | ES1M5S11026 | Moderate | P352L | 0 | 12 | 0/8 | 0/8 | 8/8 | Weak cotyledon fusion (0.41%) |
| | ES1M5S03057 | Low | S264S | 0 | 14 | 0/8 | 0/8 | 8/8 | Increased leaf serration, ectopic carpel structures |
| *HEI10* (At1g53490) | ES1M5S02042 | High | G194* | 3 | 8 | 2/16 | 9/16 | 5/16 | Reduction in chiasmata formation, reduced silique length |
| | ES1M5S10109 | Moderate | P267L | 1 | 6 | 3/8 | 4/8 | 1/8 | NT |

[a] WT and mutant refer to the number of reads supporting the presence of a wild-type and mutant sequence reads in the filtered HEM genome dataset, respectively, as compiled in the ATHEM database.
[b] NT: not tested

### Table S2: Oligonucleotide sequences used in this study

| Name | Sequence (5’-3’) | Mutations | HEM Lines |
|---|---|---|---|
| | | *CUC1*$^{L38F}$ | EH3S1B711 |
| promCUC1terfwd<br>ChCUC1Rv1 | `CAGTTGCTTGTAGAACAGGA`<br>`GGAGCTCTGCCTTTGTAAAA` | *CUC1*$^{G72E}$ | ES1M5S03067 |
| | | *CUC1*$^{E75K}$ | ES1M5S03046 |
| AtNAC1-2 RLT-Rv-<br>CUC1 | `TAGGCTTAGTGGAGACACTG`<br>`CATCGGTATGAGCAGCAGAGTT` | *CUC1*$^{G120R}$<br>*CUC1*$^{G149D}$ | ES1M5S10276<br>ES1M5S06139 |
| AtNAC1-2<br>CUC1-3' stop | `TAGGCTTAGTGGAGACACTG`<br>`TCAGAGAGTAAACGGCCACACACT` | *CUC1*$^{S177N}$<br>*CUC1*$^{P234L}$ | ES1M5S10182<br>ES1M5S01031 |
| CUC2-I2-RV<br>insituCUC2LFwd | `ACGTCACCGACTATGTCACACTCT`<br>`ATGGACATTCCGTATTACCA` | *CUC2*$^{P59L}$<br>*CUC2*$^{L62F}$ | ES1M5S10280<br>ES1M5S11077 |
| | | *CUC2*$^{G196D}$ | ES1M5S10166 |
| TAC-CUC2-TEST_R<br>insituCUC2SFwd | `AGCGGAACATCAAGCTAAGTG`<br>`GGAGGAGGAGCAACTGTG` | *CUC2*$^{S324F}$<br>*CUC2*$^{P352L}$ | ES1M5S11003<br>ES1M5S11026 |
| | | *CUC2*$^{S264S}$ | ES1M5S03057 |
| JMR83-TT2-F2<br>JMR84-TT2-R2 | `TCAAAGTGTGTATGACAGAGG`<br>`TTAGAAGCGTTCAGACAAATACA` | Splicing or intron variant<br>Splicing or intron variant<br>*TT2*$^{L234F}$ | ES1M5S02007<br>EH1S1B627<br>ES1M5S03056 |
| JMR79-TT4-F2<br>JMR97-TT4-R3 | `TATAATGGTGATGGCTGGT`<br>`AGATAGAAGGCAAGCGTTC` | *TT4*$^{W122*}$<br>*TT4*$^{T136I}$<br>*TT4*$^{G238E}$<br>*TT4*$^{V319M}$ | ES1M5S10055<br>EH1S1B670<br>ES1M5S10217<br>EH2S1B784 |
| L478<br>L479 | `GTAGAGTTGACTGGGGATTAG`<br>`GTTTTCCTGGTTCTGTTCCC` | *HEI10*$^{G194*}$<br>*HEI10*$^{P267L}$ | ES1M5S02042<br>ES1M5S10109 |
