## Supplementary Fig S1 for "A fully sequenced collection of homozygous EMS mutants for forward and reverse genetic screens in *Arabidopsis thaliana*"

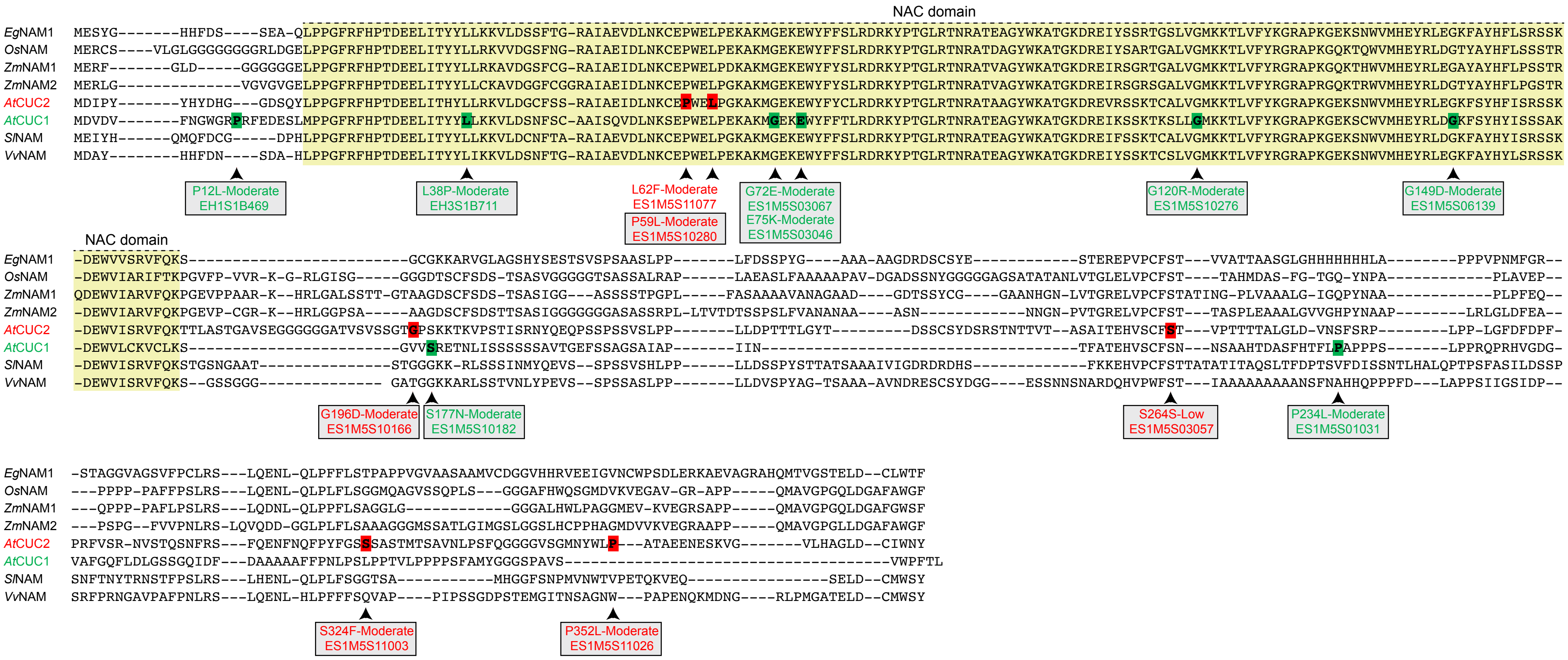


**Fig. S1: Protein sequence alignment of CUC1 and CUC2 and orthologues from *Arabidopsis thaliana*** (*At*, accession NP_188135.1, NP_200206.1, respectively), *Elaeis guineensis* (*Eg*, accession HM622270), *Oryza sativa* (*Os*, accession EAZ00836.1), *Zea mays* (*Zm*, accession CAH56057.1, CAH56058.1), *Solanum lycopersicum* (*Sl*, accession ACL14371) and *Vitis vinifera* (*Vv*, accession XP_002282655.1). Protein alignment was adapted from Adam *et al.* (2011). *At*CUC1/2 boxed residues are mutated in specific HEM lines and highlighted in green and red, respectively. The conserved NAC DNA binding domain is indicated in yellow. Boxed mutants display mutant phenotypes (see Table S1).

Adam, H. et al. Divergent expression patterns of *miR164* and *CUP-SHAPED COTYLEDON* genes in palms and other monocots: implication for the evolution of meristem function in angiosperms. Mol Biol Evol 28, 1439-1454, doi:10.1093/molbev/msq328 (2011).
